# Transcriptional profiling of *Pseudomonas aeruginosa* biofilm life cycle stages reveals dispersal-specific biomarkers

**DOI:** 10.64898/2025.12.18.695191

**Authors:** Xavier Bertran i Forga, Kathryn E. Fairfull-Smith, Jilong Qin, Makrina Totsika

## Abstract

Bacteria exhibit two lifestyles: planktonic free-floating individual cells or sessile multicellular aggregates known as biofilms. The biofilm lifecycle is characterised by three distinct stages: attachment, maturation and dispersal. Distinct adaptations occur in each stage, determining cellular behaviours such as surface attachment or synthesis and degradation of extracellular matrix components. Characterising stage-specific bacterial profiles therefore represents a valuable strategy for the development of novel antibiofilm therapies. Here, we used the model biofilm-forming bacterium *Pseudomonas aeruginosa* PAO1 to characterise the transcriptional profiles of each stage of the biofilm life cycle: attachment, biofilm maturation and spontaneous dispersal in closed cultures. We report that surface attachment was accompanied by the upregulation of genes comprising the Pil-Chp mechanosensory system, whereas biofilm maturation was characterised by the upregulation of genes involved in Pel polysaccharide synthesis, *siaD* and PA4396 diguanylate cyclases as well as *pipA, fimX* and PA5442. In contrast, dispersing cells upregulated genes responsible for the biosynthesis of alginate, rhamnolipid, and extracellular nucleases (*eddA*, *eddB*), as well as the transcriptional regulator of dispersal *amrZ*. Additionally, genes encoding the spontaneous dispersal molecule cis-2-decenoic acid (*dspS* and *dspI*), canonical phosphodiesterases (*nbdA* and *rbdA*), four non-canonical HD-GYP phosphodiesterases and seven other c-di-GMP–related enzymes were also upregulated during dispersal. Our comprehensive analysis of transcriptional changes across biofilm stages therefore provides benchmarking stage-specific transcriptional profiles for *P. aeruginosa* biofilms in closed culture systems. Furthermore, it allowed the identification of a subset of fourteen genes as transcriptional biomarkers of dispersal, which were used to build reporter plasmids as tools to determine the onset of dispersal.

**Importance:** Biofilm infections by *P. aeruginosa* are a major medical challenge due to the increased tolerance to antimicrobials displayed by bacteria living in sessile communities, which is reduced during spontaneous biofilm dispersal. Attachment, biofilm maturation and dispersal represent the main stages of a dynamic process known as the biofilm lifecycle. However, the global regulatory responses governing transitions between these stages remain understudied. Here, we combine live microscopy and biomass quantification to track the progression of *P. aeruginosa* cultures through the three main stages of the biofilm lifecycle. We show that cells from each stage recapitulate canonical, stage-specific transcriptional responses and identify a set of biomarkers associated with the onset of dispersal. These biomarkers may offer a practical tool for rapidly screening dispersal-inducing compounds, aiding in the discovery of the next generation of antibiofilm therapeutics.

## Introduction

*Pseudomonas aeruginosa* is a highly adaptable Gram-negative opportunistic pathogen, which survives in hostile environments by forming cell aggregates embedded in a protective extracellular matrix primarily composed of exopolysaccharides, extracellular DNA (eDNA) and structural proteins ^1^. These aggregates, known as biofilms, exhibit increased tolerance to desiccation and host defences compared to their planktonic counterparts ^2^. *P. aeruginosa* biofilms are associated with chronic infections, which exhibit high tolerance to a broad range of clinically employed antimicrobials ^2^.

In laboratories, *P. aeruginosa* biofilms are frequently studied using a variety of culture systems such as closed culture systems (i.e. microtiter plates) or open-flow systems (i.e. flow cells). The latter have been extensively used to identify and study the three main stages of the biofilm life cycle (attachment, biofilm maturation and dispersal), as well as distinct stage-specific molecular mechanisms ^3,4^. During attachment, cells transition from the motile to sessile lifestyle. This process is mediated by the mechanosensors Pil-Chp and Wsp which lead to the upregulation of genes involved in the synthesis of matrix components and fimbrial adhesins such as CupA ^5–7^. During biofilm maturation, the components of the matrix are progressively secreted into the extracellular space. As biofilms grow and mature, environmental cues and native intracellular signals may accumulate, such as the fatty acid dispersal signal cis-2-decenoic acid, eventually triggering spontaneous dispersal responses ^8^. This phenomenon is characterised by biofilm-residing single cells regaining motility and resuming the planktonic lifestyle, during which they escape the biofilm matrix by hydrolysing its components ^9^.

This is achieved by the upregulation of glycosyl hydrolases such as PslG and PelA for exopolysaccharide Psl and Pel degradation ^9–11^, and extracellular endonucleases EndA, EddA and EddB ^9–11^. This process is influenced by the intracellular concentration of the secondary messenger cyclic-di-GMP (c-di-GMP) ^6^, controlled by a family of proteins harbouring diguanylate cyclase domains (DGCs; possessing a GGDEF motif) and phosphodiesterase domains (PDE; possessing a HD-GYP or EAL motifs), which respectively synthesise and degrade it ^12–15^.

The transition between the stages of the biofilm life cycle is driven by distinct signalling pathways. To characterise them, transcriptomics has been applied to biofilms treated with biofilm-dispersing agents such as NO donors or carbon sources, which primarily captured immediate transcriptional changes triggered by these compounds leading to an abrupt interruption of the biofilm maturation process ^16–18^. However, the progression through the biofilm stages is a dynamic process ^19^, urging the temporal resolution of stage-specific gene expression. Therefore, we here performed RNA sequencing to compare the currently understudied transcriptional variations across cells undergoing attachment, biofilm maturation, and dispersal ^33^. Subsequently, by tracking the transcription patterns of genes throughout the three life cycle stages, we identified a subset of biomarkers for biofilm dispersal, which were used to build a collection of reporter plasmids as screening tools for the identification of the onset of dispersal.

## Results

### *P. aeruginosa* biofilms cultured in closed systems recapitulate the three defined life stages of open-flow biofilms

We previously reported that the biofilm growth kinetics of the model organism *P. aeruginosa* PAO1 in closed cultures displayed progressive accumulation of surface-attached biomass, reaching maximum biomass between 2-8 h, followed by a marked reduction ^19^. While the stages of the biofilm life cycle (attachment, maturation and dispersal) have been primarily described in open-flow cultures, these remain understudied in closed culture systems such as microtiter plates. To further characterise how these stages manifest in closed culture systems, we quantified the biofilm biomass via crystal violet staining on forming and dispersing biofilms at different time points (Fig 1A), which we combined with microscopy to capture stage-specific morphological features using microtiter plates (Fig 1B, Fig S1). Using this approach, we observed that planktonic cells initiate attachment within 2 h of inoculation (Fig 1A and B; orange), during which cells adhered to the surface as a monolayer, with a culture density (OD_600_) of ∼0.1 and a low total biomass (stained by crystal violet with OD_550_ of ∼2.0). At 4-8 h post-inoculation, cells transitioned to the biofilm maturation stage (Fig 1A and B; blue), organised in voluminous scattered clusters that grew over time and produced multicellular biofilm structures (Fig 1B, Fig S1), as observed by peak biofilm biomass staining at the 8 h time point (OD_550_ of ∼5; Fig 1A). This build-up in biomass occurred between the early (OD_600_ ∼0.3) and the late-logarithmic phase (OD_600_ ∼1.5). Beyond 8 h post-inoculation and at a higher culture density (OD_600_>1.5), biofilm biomass rapidly decreased. This phenomenon was characterised by disassembled structures leaving clearly distinguishable biofilm remnants attached to the well surface and a loss of 77% of accumulated biomass (Fig 1A and B, Fig S1; green). Collectively, our data indicate that cells grown in closed culture systems synchronously transition through distinctive phases culminating in spontaneous (endogenously induced) biofilm dispersal at defined time-points. Therefore, closed biofilm systems can effectively recapitulate the stages of the biofilm life cycle originally described in open-flow culture systems in comparatively shorter time courses ^3,20,21^.

**Figure 1.**
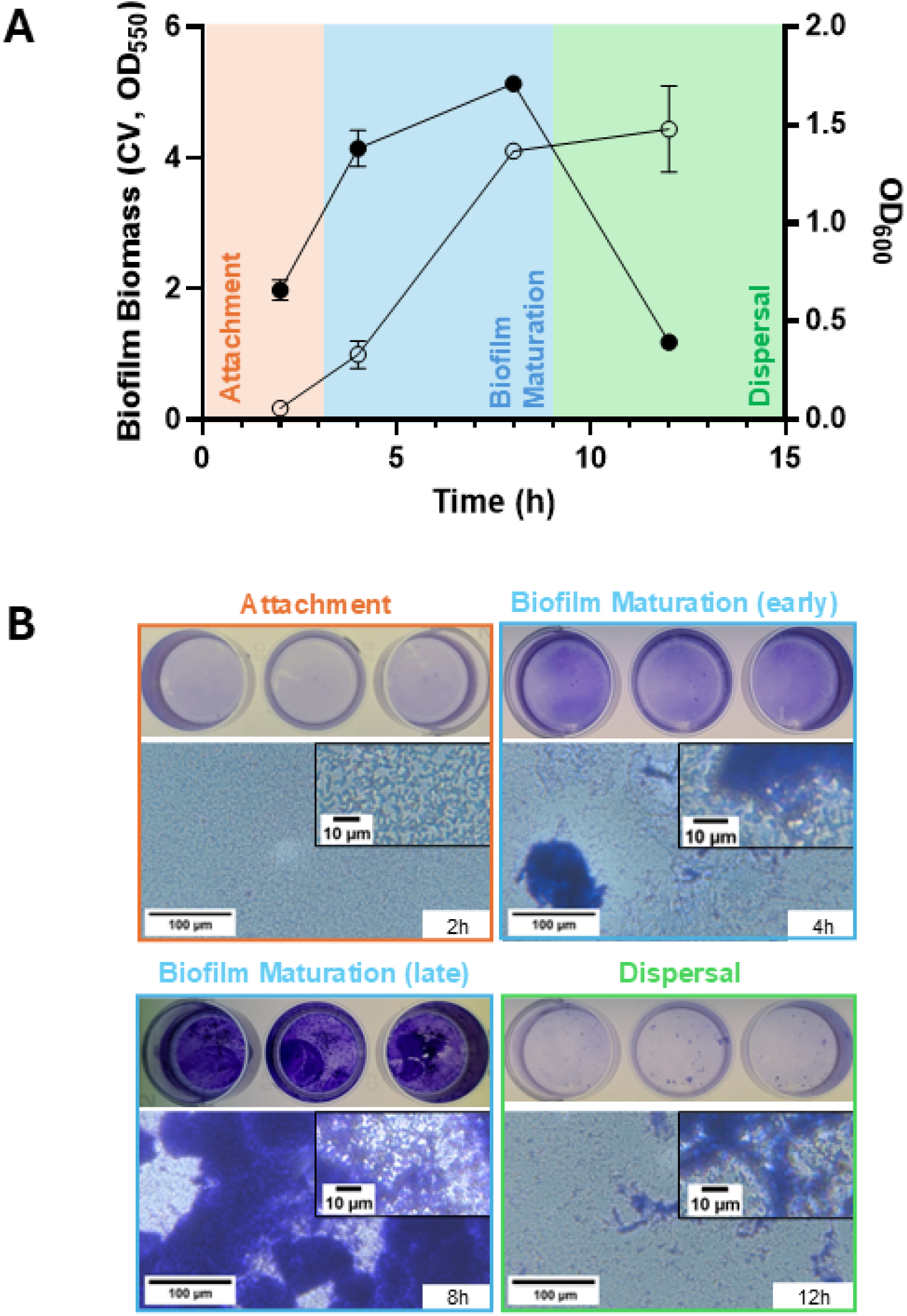
Biofilm formation and dispersal kinetics of *P. aeruginosa* PAO1. **(A)** *P. aeruginosa* PAO1 biofilm biomass and culture turbidity over 12 h. PAO1 cultures were seeded on 24-well plates and biofilm biomass was quantified at different time points by crystal violet (CV) staining (●). Culture density was simultaneously monitored by OD_600_ (○) with means ± SD shown. 3 biological replicates are represented. **(B)** Micrographs of the crystal violet-stained biofilms at each stage of the biofilm life cycle. Crystal-violet-stained microscopic images of *P. aeruginosa* biofilm cultures in 24-wells. Images are representative of at least 3 biological replicates. Orange: attachment stage. Blue: biofilm maturation stage. Green: dispersal stage. This colour coding scheme is used throughout the article to differentiate the three biofilm stages.

### Transcriptional profiling of *P. aeruginosa* biofilm cultures displays canonical pathways of attachment, maturation and dispersal

While *P. aeruginosa* cells grown in microtiter plates undergo the three canonical stages of the biofilm life cycle, RNA recovered from biofilm cells cultured in microtiter plates is typically insufficient in quantity to support downstream RNA sequencing (RNA-seq). Therefore, we replicated the biofilm kinetics in tissue culture flasks as a larger-volume closed culture system, monitoring biofilm cell density in addition to total biomass via staining with crystal violet. Biofilm kinetics displayed by *P. aeruginosa* PAO1 in tissue culture flasks mimicked the temporal pattern of microtiter plates in total biomass and microscopically (Fig S1), showing attachment, biofilm maturation and dispersal within the same time frames.

Using the flask culture system, we performed RNA sequencing of *P. aeruginosa* during attachment, early biofilm maturation and dispersal to characterise stage-specific transcriptional profiles (Fig 2A). This study design enabled temporal resolution of both global and individual gene transcription variation across the biofilm life cycle stages. Accordingly, biofilm maturation was used as the central reference point, as it represents the intermediate stage between attachment and dispersal, (Fig 2C). Differential transcription analysis using a negative binomial test (*P-*value ≤ 0.01; Log_2_ fold-change ≥ |1|) identified a total of 3361 genes with significant transcriptional changes across all three comparisons: i) Attachment vs. Biofilm Maturation; ii) Biofilm Maturation vs. Dispersal (Planktonic); and iii) Attachment vs. Dispersal (Table S1). Cells collected from the liquid phase and from biofilms during at the dispersal stage showed few transcriptional differences, with only 83 differentially regulated genes (76 upregulated and 7 downregulated) between the two states, and with low fold changes (Table S1). This suggested that cells at the dispersal stage measured in this study synchronously altered their transcriptomic profiles regardless of remaining surface-associated or becoming free-swimming. Due to the planktonic phase of dispersal containing larger transcriptional variations relative to other stage transitions (Fig 2B), we used it for all subsequent analyses. Principal component analysis (PCA) further confirmed that cells at each stage formed distinct clusters, indicating discrete transcriptional profiles were associated with attachment, maturation, and dispersal (Fig 2B).

**Figure 2.**
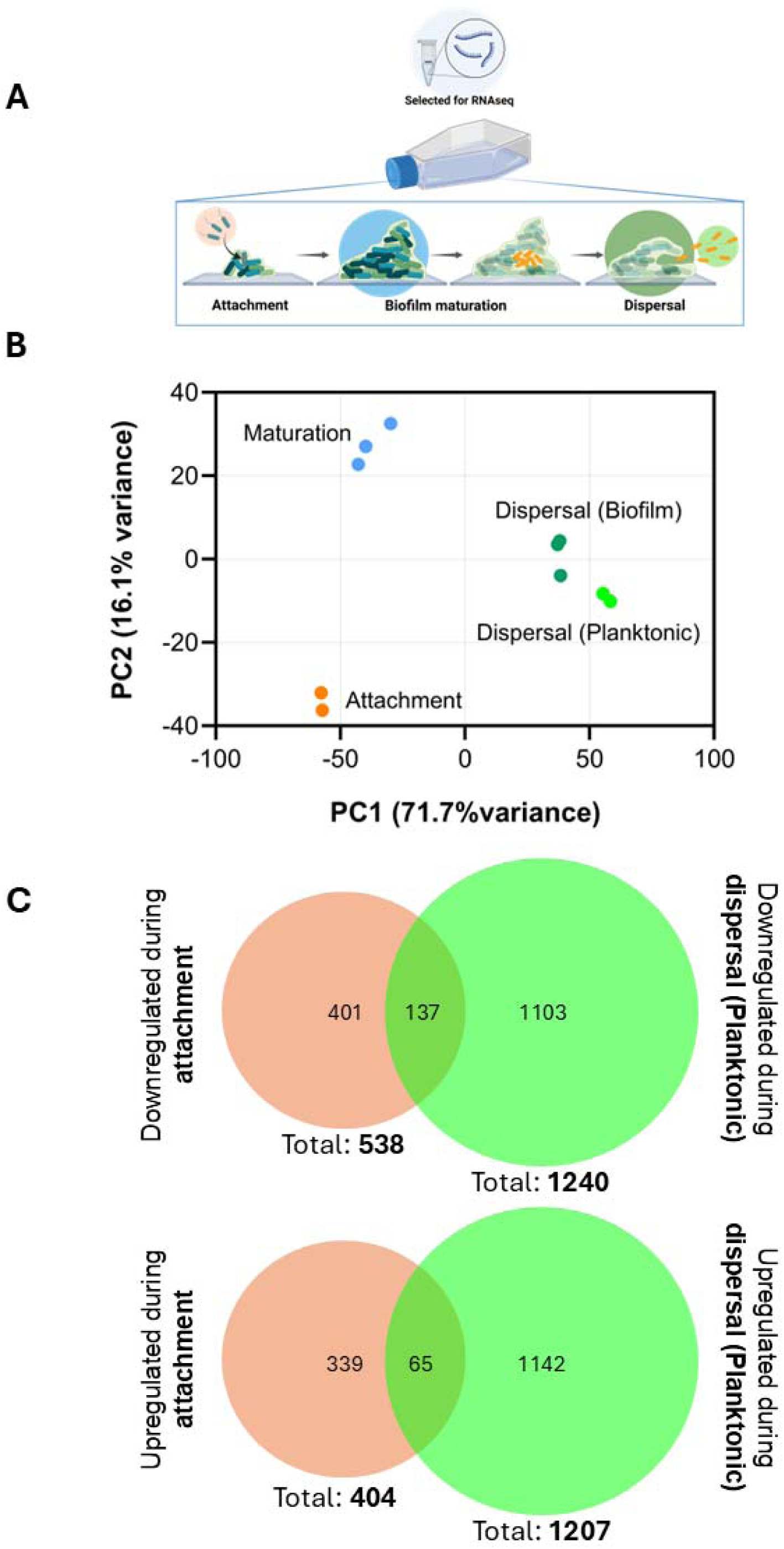
Transcriptional profiles of different stages of the biofilm life cycle. **(A)** Schematic depicting RNA sample collection for sequencing. During attachment, the free-swimming phase was collected to represent cells rapidly transitioning from motile to sessile. During biofilm maturation, surface-attached cells were collected, and cells from both phases were collected during the dispersal stage. **(B)** Principal component analysis of samples collected at distinct stages of the biofilm life cycle (attachment – orange; biofilm maturation – blue; the planktonic phase of biofilm dispersal – light green; and the biofilm phase of biofilm dispersal – dark green). Each biological replicate of a group is represented as a dot of the same colour. The normalised reads of all genes were then extracted for each sample and transformed into PC loadings with ClustVis ^22^. Values for PC1 and PC2, accounting for the largest variance percentage are represented in the plot. **(C)** Venn diagrams showing differentially transcribed genes in cells collected from the liquid phase undergoing attachment and dispersal (relative to biofilm maturation). Downregulated (top) and upregulated (bottom) genes at each stage relative to biofilm maturation are represented separately. Overlapping areas indicate genes with consistent transcriptional changes relative to biofilm maturation stage.

Building on these global transcriptional variations and to further validate our datasets, we mined our RNA-seq data for previously reported stage-specific changes in the transcription of genes previously associated with attachment, biofilm formation or dispersal. During attachment and extending into biofilm maturation, we observed the upregulation of the Pil-Chp surface-sensing complex known to mediate attachment ^7^, whereas no upregulation of the complementary Wsp system was noted. Notably, transcription of *pilA, pilB, pilD* and *pilGHIJK-chpABCDE,* encoding type IV pili structural and signal transduction proteins, and of *cyaB* and *vfr,* which encode an adenylate cyclase and its associated cyclic-AMP-controlled transcriptional regulator Vfr, was upregulated between 2.00- and 7.37-fold relative to biofilm dispersal.

Biofilm maturation involves synthesis pathways for extracellular polysaccharides and eDNA, which are the main components of the biofilm matrix ^1^. During biofilm maturation, we observed the upregulation of the exopolysaccharide Pel synthesis operon *pelBCDEF* but not *pelG* relative to attachment (2.22 – 2.26-fold) and dispersal stages (2.16 – 2.99-fold) (Fig 3A, Table S1) ^23^. Additionally, the alternative respiration process of denitrification is known to be active during biofilm maturation due to cell layering inside the biofilm restricting oxygen availability ^24^. Accordingly, we found biofilm cells to upregulate the transcription of the nitrite reductase complex (*nirSMCFDLG*) by up to 10-fold relative to attachment.

**Figure 3.**
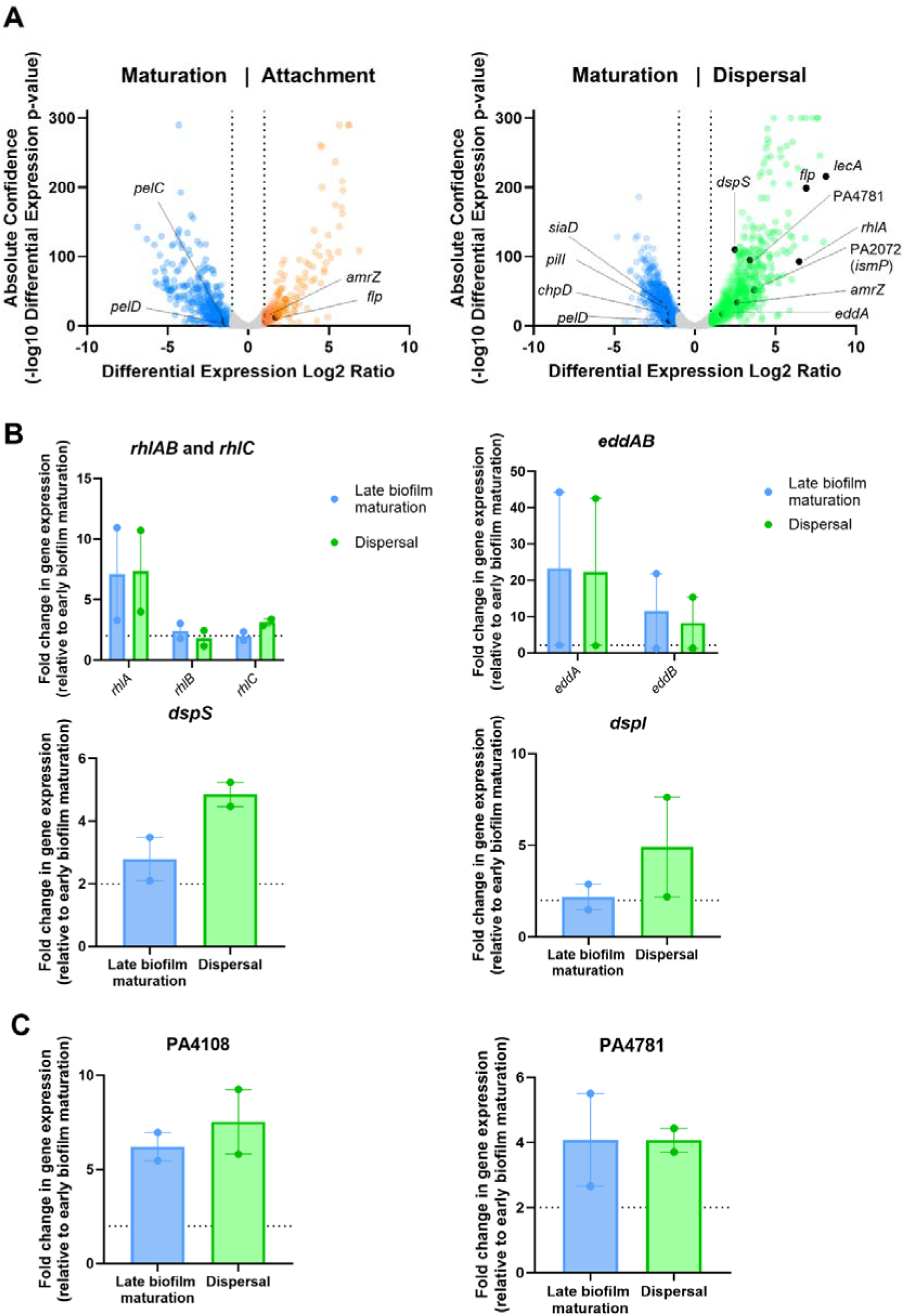
Genes involved in dispersal mechanisms are upregulated during late biofilm maturation and dispersal compared to early biofilm maturation. (**A)** Comparison of the transcriptomes of planktonic cells during attachment or dispersal to that of sessile cells during biofilm maturation. Genes containing Log2-fold change transcription values >|1| were highlighted in biofilm stage specific colours. (**B and C)** RT-qPCR analysis of canonical dispersal genes (*rhlA, rhlB, rhlC, eddA, eddB, dpsS* and *dspI*) and the PDEs PA4108 and PA4781, which were identified as putatively involved in dispersal. RNA samples were extracted at different time points with Means ± SEM shown.

Planktonic cells at the biofilm dispersal stage, relative to attachment and biofilm maturation stages, showed an increased transcription of genes involved in eDNA hydrolysis (*eddA,* 3.28-fold; and *eddB,* 2.95-fold; but not *endA*) (Fig 3A, Table S1), suggesting the remodelling of the extracellular matrix to allow the escape of planktonic cells. We also observed a pronounced upregulation of quorum-sensing–regulated genes previously shown to drive biofilm dispersal. *rhlA*, *rhlB*, and *rhlC*, which encode enzymes for the biosynthesis of the rhamnolipid biosurfactant implicated in quorum-sensing–mediated dispersal ^39–42^were upregulated by 88-, 45- and 12-fold, respectively (Fig 3A, Table S1) ^25–28^. Another quorum-sensing pathway activated during dispersal involved the fatty acid signal cis-2-decenoic acid ^27,29^. This fatty acid signal is produced and sensed by the proteins encoded by *dspI* and *dspS*, which were upregulated by 2.04- and 5.59-fold, respectively, during dispersal relative to the biofilm maturation stage (Fig 3A, Table S1). Interestingly, genes associated with biofilm maturation were also found upregulated during the dispersal stage. Adhesion factors, such as lectins *lecA* and *lecB* were highly upregulated during dispersal by 279-fold and 115-fold, respectively, relative to the biofilm maturation stage. Similarly, *flp*, which encodes the main subunit of the type IVb pili adhesin was upregulated by 120-fold, together with several alginate biosynthesis genes (*alg8*, 2.26-fold*; algX,* 2.16-fold*; algJ*, 2.39-fold; *algA,* 2.74-fold) and their associated regulators *algU* (2.49-fold)*, algR* (5.76-fold) and *amrZ* (6.27-fold) (Fig 3A, Table S1). These findings were consistent with previous studies showing that dispersed cells had significantly increased functional adhesion relative to planktonic cells ^30^. Therefore, our RNA-seq data captured transcriptional changes associated with previously documented attachment, biofilm maturation and dispersal responses, further underscoring the consistency of our dataset with established biofilm regulatory pathways and the robustness of our analytical methods.

To further validate the upregulation of these known dispersal pathways, we extracted RNA from biofilms at 4 h (early maturation), 8 h (late maturation), and planktonic cells collected at 12 h (dispersal) and measured the transcript levels of representative genes involved in rhamnolipid synthesis (*rhlAB, rhlC*), eDNA degradation (*eddA, eddB*), and cis-2-decenoic acid signalling (*dspI, dspS*) by RT-qPCR (Fig 3B). Relative to maturing biofilms, all genes were upregulated in mature and dispersed biofilms. These findings further validate our RNA-seq data and indicate that mature biofilms may begin to transcriptionally activate dispersal pathways before the onset of active detachment.

### Specific c-di-GMP enzymes characterise the biofilm maturation and dispersal transcriptomes

Cyclic di-GMP (c-di-GMP) is a key signalling molecule governing the transition from biofilm maturation to dispersal. Considering that the roles of enzymes responsible for its synthesis and breakdown are stage-specific, we also analysed our transcriptomic data to identify genes encoding enzymes with predicted PDE (EAL or HD-GYP) or DGC (GGDEF) domains (Table 1). Expectedly, genes encoding proteins essential for biofilm development (*siaD,* 3.73-fold and *pipA,* 2.08-fold) were upregulated during biofilm maturation ^31–34^. Genes implicated in the formation of large microcolonies ^32^, including PA4396 (2.08-fold), *fimX* (2.77-fold), and PA5442 (2.36-fold) were also significantly upregulated. In contrast, we observed an upregulation of a combination of PDEs and DGCs in cells at the dispersal stage, including sensory transducers of dispersal cues *nbdA* (7.56-fold) and *nicD* (6.73-fold) ^35–37^. Additionally, upregulation of *ismP* (12.99-fold) was noted, which encodes an iron sensor that under low iron conditions hijacks and inhibits a cognate DGC ImcA ^38^.

**Table 1.**
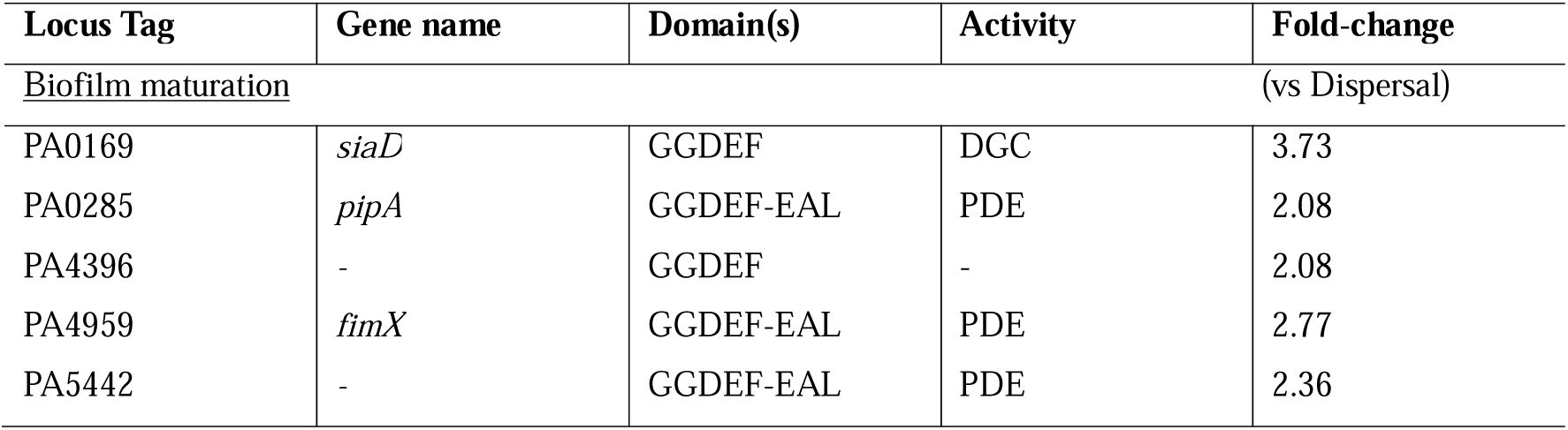
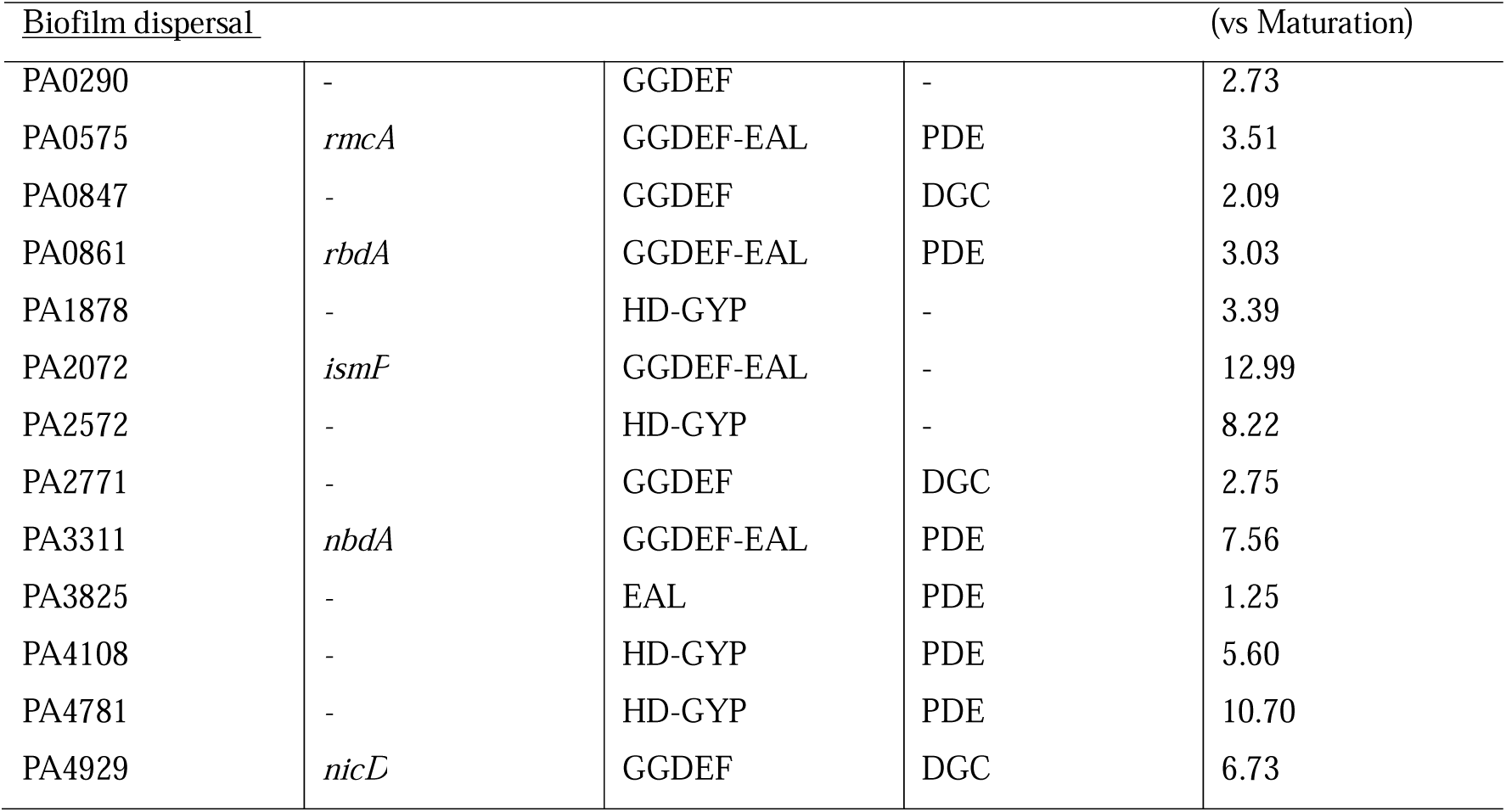
Differentially regulated genes encoding c-di-GMP-modifying enzymes during biofilm maturation and dispersal.

Interestingly, dispersed cells upregulated all four genes encoding proteins with HD-GYP phosphodiesterase domains present in the *P. aeruginosa* PAO1 genome (PA1878, PA2572, PA4108 and PA4781) (Table 1). Two of these, PA4108 and PA4781 were previously reported to have PDE activity and to be necessary for swarming and the production of virulence factors ^39,40^. We further validated the transcriptional changes of these two genes using RT-qPCR (Figure 3C), confirming an increase in expression of 7.5-fold and 4-fold, respectively, during dispersal relative to maturing biofilms at 4 h. Moreover, the transcription of both genes was also upregulated in mature biofilms (6.2- and 4-fold; Figure 3C), indicating a similar pattern of transcriptional priming prior to dispersal. Altogether, our results indicate that canonical stage-specific c-di-GMP modulation responses are accurately reflected in our transcriptomic dataset and suggest a potential undescribed role of HD-GYP-containing PDEs in mediating biofilm dispersal.

### Transcriptional biomarkers identify the onset of biofilm dispersal

Since the obtained transcriptomic profiles recapitulated canonical pathways associated with attachment, maturation, and dispersal, our datasets are uniquely well suited to identify gene changes associated with the dispersal phase of *P. aeruginosa* and serve as transcriptional biomarkers of biofilm dispersal. We reasoned that genes directly involved in biofilm dispersal would exhibit a distinct temporal pattern of transcription such as an inverse regulation between the biofilm maturation and dispersal stages relative to the initial attachment stage. Thus, pro-dispersal genes would be repressed during maturation but strongly upregulated during dispersal, or vice versa. To identify candidate biomarkers, we applied these selection criteria and identified 14 upregulated and 8 downregulated genes that could potentially serve as dispersal biomarkers (Table 2).

**Table 2.** Genes differentially regulated at the biofilm maturation and dispersal stage compared to the attachment stage.

| Gene | Fold change in<br>Biofilm<br>maturation <sup>a</sup> | Fold change in<br>Biofilm<br>dispersal <sup>a</sup> | Gene product |
| --- | --- | --- | --- |
| Downregulated at 4h and upregulated at 8h |  |  |  |
| PA0111 | -2.01 | 12.21 | hypothetical protein |
| *cheR2* | -2.22 | 2.10 | chemotaxis protein methyltransferase |
| PA0743 | -2.58 | 3.12 | NAD-dependent L-serine dehydrogenase |
| PA1353 | -2.36 | 3.32 | hypothetical protein |
| *pqqA* | -5.10 | 2.55 | coenzyme PQQ synthesis protein A |
| PA2588 | -2.03 | 3.01 | transcriptional regulator |
| *amrZ* | -2.30 | 2.55 | alginate and motility regulator Z |
| *tadA* | -2.17 | 13.64 | ATPase TadA |
| *rcpA* | -2.13 | 12.47 | type II/III secretion system protein |
| *rcpC* | -2.07 | 13.83 | hypothetical protein |
| *flp* | -3.27 | 34.54 | type IVb pilin Flp |
| PA4523 | -2.31 | 3.41 | hypothetical protein |
| *cupE1* | -2.11 | 9.58 | fimbrial subunit CupE1 |
| *cupE2* | -2.04 | 3.95 | fimbrial subunit CupE2 |
| Upregulated at 4h and downregulated at 8h |  |  |  |
| PA0485 | 2.93 | - 2.91 | hypothetical protein |
| PA1170 | 2.43 | -4.76 | hypothetical protein |
| *metH* | 2.45 | -2.13 | B12-dependent methionine synthase |
| *ilvH* | 3.53 | -2.35 | acetolactate synthase small subunit |
| *ilvI* | 4.69 | -2.57 | acetolactate synthase 3 catalytic subunit |
| *cysN* | 2.1 | -2.62 | bifunctional sulfate adenylyltransferase<br>subunit1/adenylylsulfate kinase |
| PA4889 | 2.06 | -14.92 | oxidoreductase |
| PA4907 | 2.41 | -2.27 | short-chain dehydrogenase |

<sup>a</sup> Relative to samples collected during attachment.

**Table 3.**
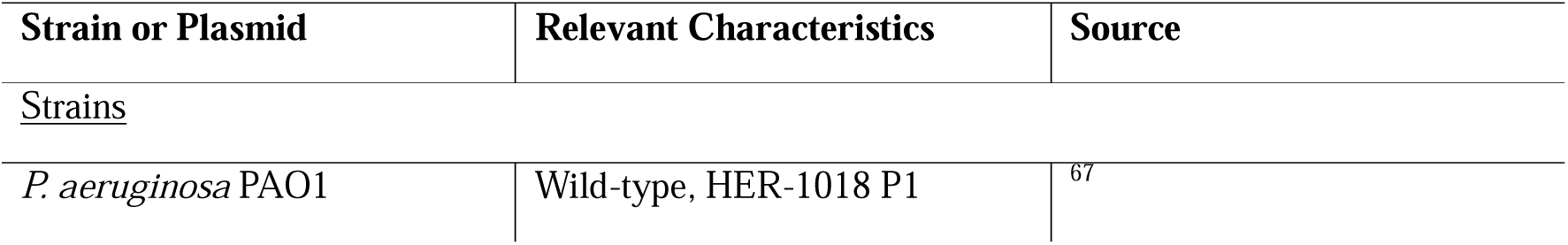
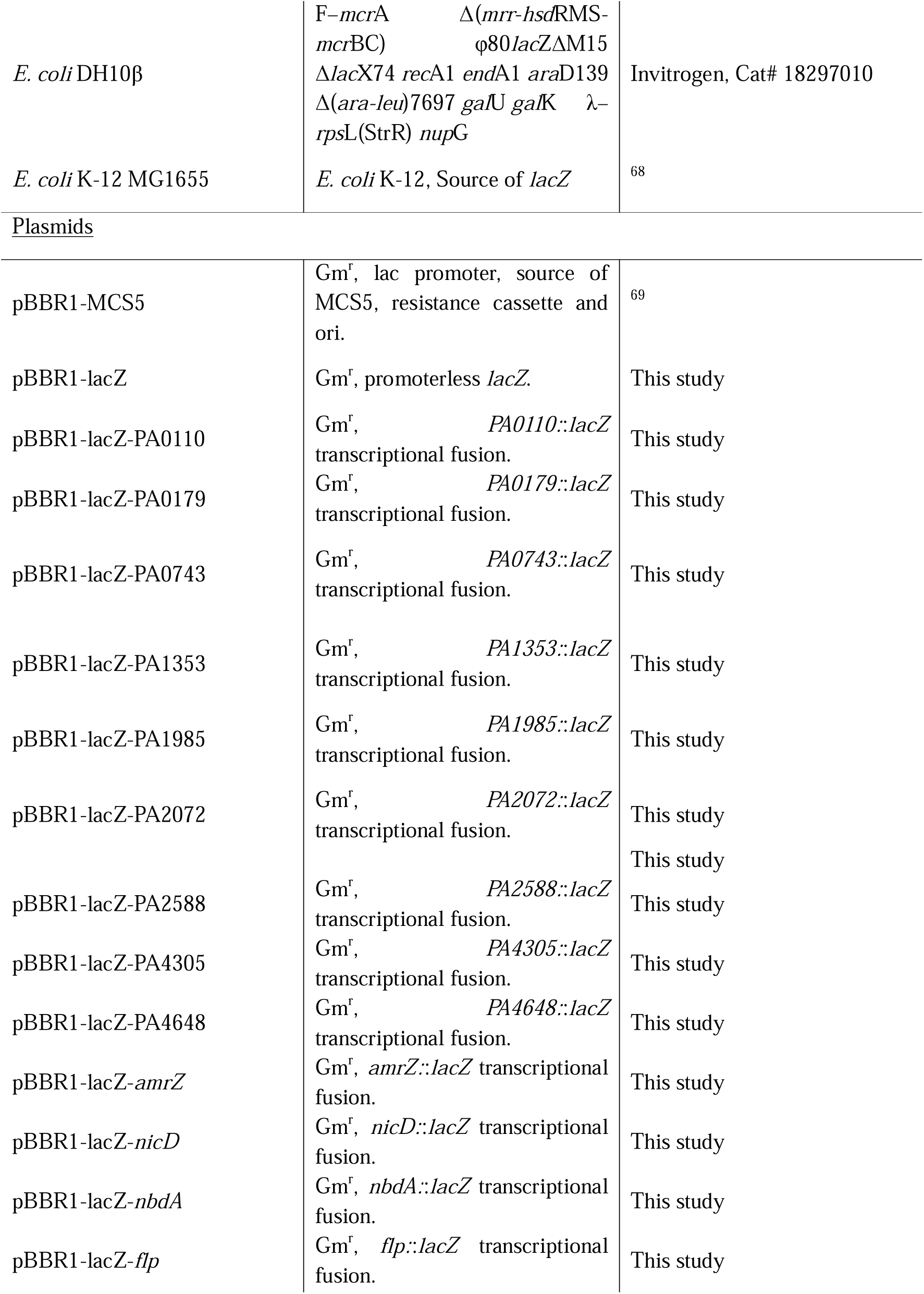
Bacterial strains and plasmids used in this study.

Among the upregulated genes, *cheR2* (2.1-fold, Table 2) encodes a methyltransferase component of the Che2 chemotaxis pathway, which interacts with the heme-containing oxygen sensor McbP/Aer2 and the coupling protein CheW2 to modulate virulence through a cryptic phosphorylation cascade ^41,42^. Another notable gene, *pqqA* (2.55-fold; Table 2), is part of the *pqqABCDE* operon responsible for the biosynthesis of pyrroloquinoline quinone (PQQ), a redox cofactor for the quinoprotein ethanol dehydrogenases ExaA and ExaBC ^43^. In addition, several genes under the transcriptional control of the two-component system PprAB were also upregulated. These include *tadA*, *rcpA*, *rcpC*, and *flp* (13,66-, 12.47-, 13.83- and 34.54-fold; Table 2) which are part of a contiguous cluster spanning PA4297 (*tadG*) to PA4306 (*flp*). This locus encodes the machinery for the synthesis and secretion of Flp type IVb pili ^44^. Notably, two of the dispersal biomarkers *amrZ* and PA2588 (*cdpR*) have known roles in modulating biofilm physiology. *amrZ* encodes a recognised transcriptional regulator of dispersal and *cdpR* is a quorum sensing and virulence transcriptional regulator ^45–47^.

Several uncharacterised genes, including PA0111, PA0743 and PA1353 were also identified as upregulated biomarkers of biofilm dispersal, many of which appear to be associated with redox processes. While no experimental data are available for PA0111, bioinformatic predictions suggest that the directly upstream gene PA0110 encodes a protein putatively related to a mitochondrial cytochrome *c* oxidase. Additionally, PA0112 and PA0113 are predicted to encode a heme A synthase and a protoheme IX farnesyltransferase, respectively, implicating these loci in cytochrome maturation or electron transport. Similarly, PA0743, which encodes an L-serine dehydrogenase, was also upregulated, potentially reflecting altered amino acid catabolism related to redox homeostasis during biofilm dispersal. No functional predictions are currently available for PA1353, which was also part of the identified dispersal biomarkers.

To evaluate the wider use of these biomarkers beyond the reference PAO1 strain, we investigated gene presence across 1301 *P. aeruginosa* genomes. Indeed, we identified that across a diverse range of genomic backgrounds, all fourteen transcriptional biomarkers of dispersal are present in ∼100% of complete *P. aeruginosa* genomes available in GenBank, with only *flp* showing an average pairwise identity <99% (Table S2). We restricted our biomarker screen to upregulated transcripts, as elevated expression levels are more consistently detected and validated across analytical platforms. Due to the similarities in transcriptomic profiles between planktonic and biofilm cells collected during dispersal, we hypothesised that biofilm cells initiate the expression of dispersal biomarkers prior to phenotypically manifesting the dispersal response in culture. Therefore, we collected samples from cells undergoing late biofilm maturation and dispersal and validated these biomarker genes by RT-qPCR (Fig 4). Our results confirmed the upregulation of all fourteen genes during dispersal. Furthermore, all genes but four (PA0743, PA1353, *pqqA*, and PA2588) were also significantly upregulated during late biofilm maturation (Fig 4). Genes PA0111, *cheR2*, *tadA*, *rcpC*, *flp*, and PA4523 showed a pattern where transcript abundance increased in mature biofilms (2.5- to 23-fold) and peaked during dispersal (4.6- to 117-fold relative to early biofilms) (Fig 4). In contrast, the transcription of *cupE1* and *cupE2* fimbrial genes peaked in mature biofilms (54.41- and 10.1-fold) but sharply declined upon dispersal (25.55- and 5.9-fold) (Fig 4). This transient transcriptional profile suggests that CupE fimbriae genes are likely characteristic of mature biofilms. In contrast, transcription of PA0111, *cheR2, tadA, rcpC, flp* and PA4523 peaked at dispersal (Fig 4). Altogether, these biofilm dispersal biomarkers exhibited strong reproducibility, and their increased transcription prior to the onset of dispersal marks biofilm cells undergoing late biofilm maturation that are ready for an incoming stage transition.

**Figure 4.**
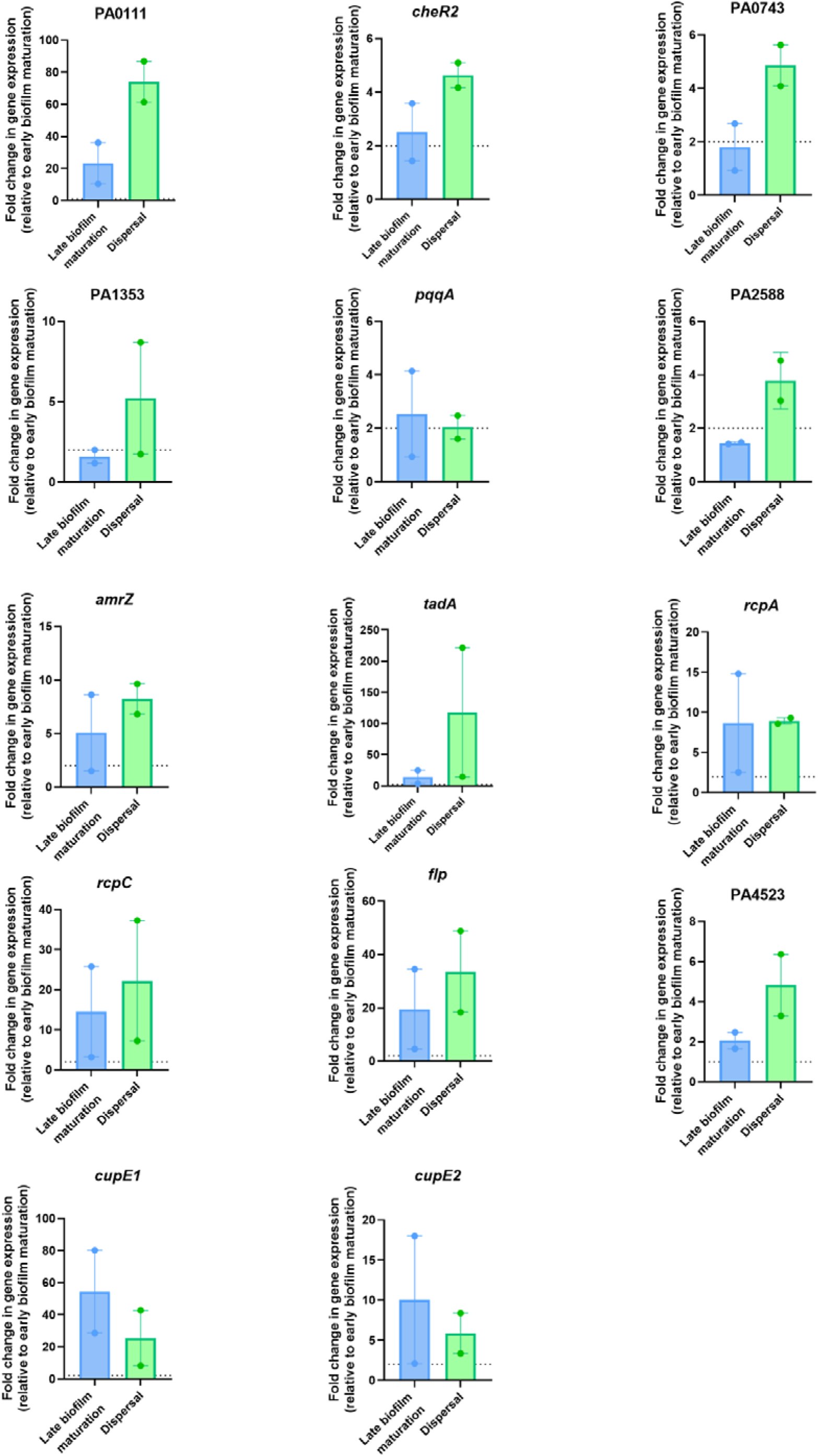
Validation of the transcriptional biomarkers of biofilm dispersal. Differences in transcriptional regulation measured by RT-qPCR for the indicated genes from total RNA samples extracted at 4 h (early biofilm maturation), 8 h (late biofilm maturation) and 12 h (dispersal). Fold-changes at 8 h and 12 h are relative to 4 h. Means ± SEM are shown. Two technical replicates are represented. Statistical differences between groups were calculated using Student *t-*tests (*, *P*-value <0.05).

### Reporter plasmids as a low-cost rapid tool for monitoring biofilm dispersal

While RT-qPCR provides a direct and sensitive method to quantify transcriptional differences, we constructed a more scalable reporter system using plasmids carrying *lacZ* transcriptional fusions to the predicted promoter regions of each biomarker, excluding co-operonic genes (*rcpC* also represents *rcpA* and *tadA*, and *cupE1* also represents *cupE2*). To evaluate this approach, *P. aeruginosa* PAO1 strains harbouring each reporter plasmid were cultured to early biofilm maturation (4 h) and dispersal (12 h). β-galactosidase activity was significantly increased for 9 of 11 reporters, with only *cheR2* and *flp* showing elevated levels of enzymatic activity at both early biofilm maturation and dispersal (Fig 5). In contrast, reporters containing PA0110, PA0743, PA1353, *pqqA*, PA2588, *amrZ*, *rcpC/rcpA*, PA4523, and *cupE1/cupE2* showed significantly increased enzymatic activity during dispersal relative to early biofilm maturation (Fig 5). Negative enzymatic activity values for *amrZ* at 4 h indicate a very low presence of LacZ, where background noise levels (OD_550_) are similar to total recorded activity (OD_420_; see Methods). Collectively, these results demonstrate that *lacZ* transcriptional reporter plasmids provide a low-cost and rapid detection method for the transcriptional activation of the identified biomarkers of dispersal and further validate these genes as robust biomarkers of the onset of dispersal.

**Figure 5.**
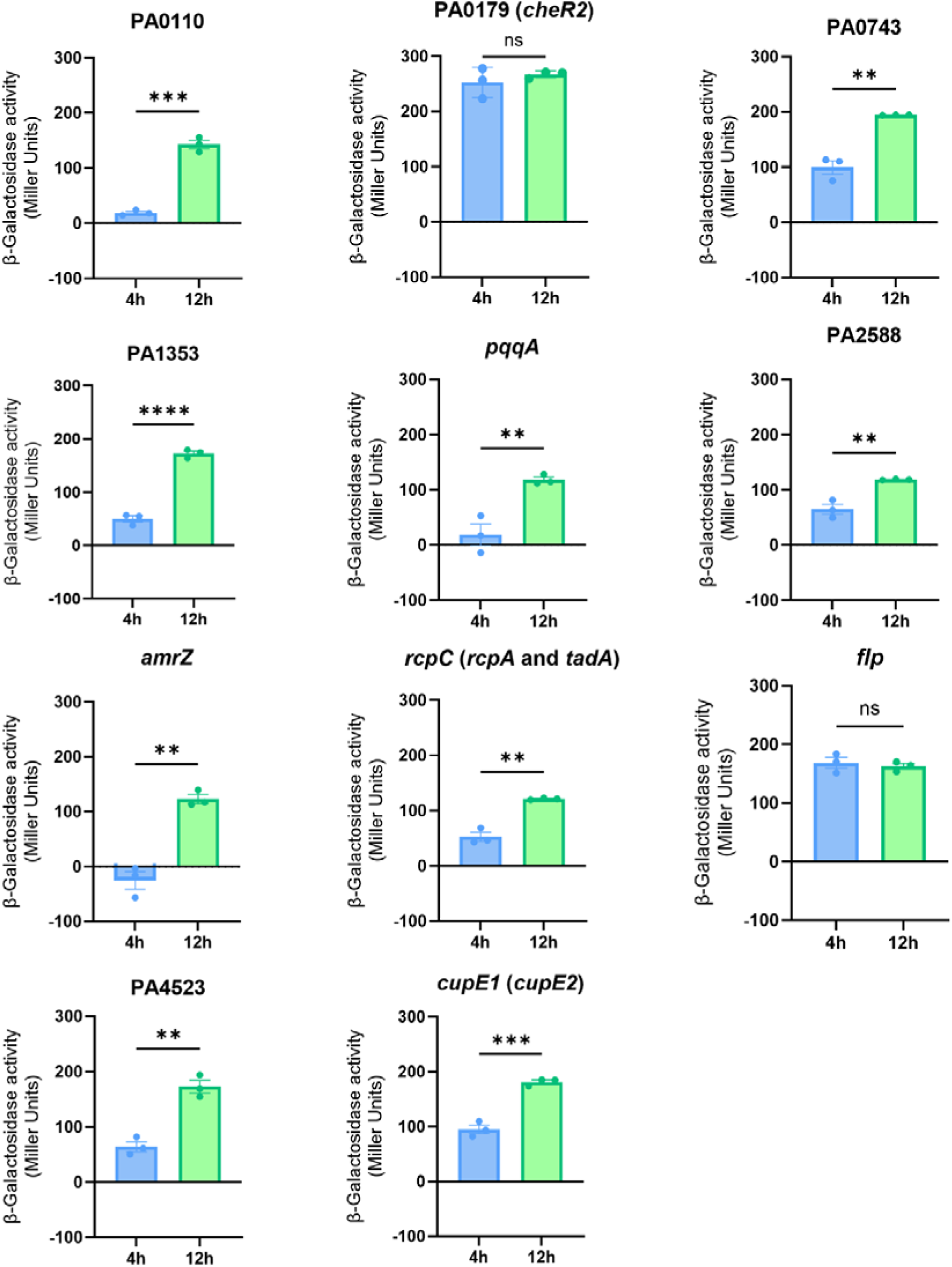
LacZ reporter plasmids indicate the upregulation of the identified biomarkers during dispersal. Differences in transcriptional regulation of biomarkers of dispersal measured by enzymatic activity of β-galactosidase in *P. aeruginosa* PAO1 in biofilm samples extracted at 4 h (early biofilm maturation) and 12 h (dispersal). Means ± SEM are shown. Three technical replicates are represented. Statistical differences between groups were calculated using Student’s *t-*tests (**, *P*- value <0.01; ***, *P*-value <0.001; ****, *P*-value <0.0001). One representative experiment is shown from two independently performed assays.

## Discussion

The study of transcriptional changes between each biofilm stage provides valuable insights into the key cellular signalling pathways underlying their transition between sessile and motile lifestyles. Here, we characterised the transcriptional profiles of cells undergoing attachment, biofilm maturation and dispersal, and identified 23 biomarkers characterising the dispersal response, 14 of which comprising a subset upregulated upon the initiation of dispersal.

Cells undergoing attachment sense and respond to surface contacts through mechanosensory systems such as the Pil-Chp machinery, which relies on type IV pili-mediated twitching motility ^48,49^. In our study, we report an upregulation of type IV pili biogenesis genes (*pilA, pilB* and *pilD*) and of the mechanosensory system Pil-Chp (*pilGHIJK-chpABCDE*) during the attachment and biofilm maturation stages compared to dispersing cells. These were accompanied by the upregulation of *fimX*, which encodes a putative phosphodiesterase essential for the assembly of type IV pili, and of *cupA1* encoding the main subunit of the CupA adhesin ^50^. Our findings therefore align with previous studies indicating that type IV pili are indispensable to surface-attachment in microplates ^51^, and are consistent with prior reports that surface-sensing promotes the transcriptional upregulation of genes involved in type IV pilus biogenesis ^52^.

The intracellular messenger c-di-GMP acts as a master regulator of the bacterial lifestyle, with its accumulation leading to biofilm formation ^9^. Our dataset revealed the upregulation of several stage-specific genes encoding proteins with validated or hypothetical c-di-GMP modulating activity. Specifically, *siaD* was the most upregulated during biofilm maturation, which encodes a DGC involved in early microcolony formation and adhesion ^48,53^. In contrast, biofilm dispersal is associated with increased PDE activity and decreasing intracellular c-di-GMP ^54^. This was reflected in our dataset, where 6 genes encoding functionally active PDEs were upregulated. Genes *rbdA* and *nbdA* encode enzymes with well-described roles in biofilm dispersal. Both have been reported to modulate flagellar motility, with RbdA being a key component of the BdlA-regulated core dispersal response ^55,56^. Moreover, four upregulated genes (PA1878, PA2572, PA4108 and PA4781) encode PDEs carrying the HD-GYP motif ^57^. While the role of these genes in the biofilm dispersal stage is unclear, it has been reported that disruption of PA4108 and PA4781 resulted in higher intracellular c-di-GMP and decreased swarming activity, whereas PA2572 modulated swarming motility ^57,58^. We here validated by RNA-seq and RT-qPCR that both PA4108 and PA4781 are strongly upregulated during late biofilm maturation and dispersal, which suggests a potential role in initiating dispersal responses. Additionally, recent reports showed that deletion of *cdpR* led to increased biofilm biomass and decreased swarming motility ^47^. Considering that the upregulation of *cdpR* is strongly controlled by the quorum sensing regulator RsaL ^46^, the stage-specific transcriptional activation of *cdpR* would therefore suggest that dispersal in closed systems is mediated by a global quorum sensing-mediated swarming response.

The synthesis of the rhamnolipid biosurfactant is important to the quorum sensing-mediated swarming motility and dispersal ^26^. In closed culture systems, rhamnolipid was proposed to drive biofilm dispersal ^59^. Here we report that the rhamnolipid and swarming modulator PA2572 together with several rhamnolipid-associated genes were highly upregulated during dispersal. Moreover, four of the identified PDEs upregulated during dispersal, including PA4108, PA4781, *nbdA* and *rbdA,* were previously shown to influence the synthesis of rhamnolipid leading to loss of biofilm biomass ^20,36,58^. Altogether, these data suggest biofilm dispersal in closed systems may be driven by rhamnolipids.

Among the upregulated biomarkers, *amrZ* was reported as a global regulator controlling the upregulation of genes related to alginate biosynthesis and Type IV pili twitching motility while downregulating Pel synthesis, therefore driving biofilm dispersal ^45,60^. In contrast, *tadA*, *rcpA*, *rcpC*, *flp, cupE1* and *cupE2*, which are regulated under the PprAB two-component system ^61,62^, were reported to mediate surface attachment. Our data therefore support previous studies indicating that dispersing cells exhibit a hyperadhesive potential that primes them to re-attach to surfaces ^30^. Additionally, we identified PA0110, *cheR2*, *pqqA,* PA0743, PA1353 and PA4523 to be uniquely associated with the dispersal stage, yet their specific roles remain to be defined. Their marked upregulation during dispersal warrants further investigation.

Understanding biofilm dispersal may represent a critical step toward combating chronic biofilm-associated infections. As dispersing cells become resensitised to antimicrobial treatment once they are released from the protective biofilm matrix, the underlying transcriptional changes occurring during biofilm dispersal hold promise for unveiling the next generation of antibiofilm treatments and adjuvants. We here report the transcriptional profiles of the *P. aeruginosa* biofilm life cycle, including attachment, biofilm maturation and dispersal stages in closed systems, which can be used as benchmarks for future studies. Additionally, we propose a set of biofilm dispersal biomarkers that are robustly activated at the onset of biofilm dispersal. As the upregulation of these biomarkers was consistently reproduced by transcriptomic and RT-qPCR analyses, we generated gene fusion reporters that enable the rapid monitoring of promoter activities *in vivo,* as well as being adaptable to high-throughput (i.e. microplate-based) platforms. Therefore, these reporter plasmids present with potential application in high throughput screens supporting the discovery and development of novel dispersal-inducing antibiofilm therapeutics.

## Materials and Methods

### Strains, media and culture conditions

*Pseudomonas aeruginosa* PAO1 wild-type (WT). Cultures were routinely grown overnight in LB (lysogeny broth) media at 37°C, 200rpm before incubation in fresh M9 media (9 mM NaCl, 22 mM KH_2_PO_4_, 48 mM Na_2_HPO_4_, 19 mM NH_4_Cl and 2 mM MgSO_4_, 100 µM CaCl_2_, 0.4% glucose, pH 7.0) for each experiment. Where appropriate, cultures were supplemented with 10µg/ml of gentamicin.

### Biofilm kinetics and microscopy

Biofilms were grown on 24-well plates or tissue culture flasks (Nunc, ThermoFisher Scientific) by inoculating 10^7^ colony forming units (CFU)/ml of overnight bacterial cultures for 2 h, 4 h, 8 h and 12h diluted in M9 media. 1 ml of culture was inoculated in each well of microtiter plates and 10 ml were used in tissue culture flasks. Cultures were incubated at 37 °C and 180 rpm (microtiter plates) or 70 rpm (tissue culture flasks). Shaking speed was adjusted to prevent splashing. At the indicated time points, supernatants were removed, and wells were delicately washed with 1ml phosphate buffered saline (PBS, Gibco) to remove planktonic cells. Microscopy (PrimoVert, Zeiss) was used to confirm the removal of planktonic cells and preservation of biofilm structures. Remaining biofilm cells were stained with 0.1% (w/v) crystal violet dissolved in 6.25% (v/v) methanol. Stained biofilms were washed twice with PBS and solubilised with ethanol. Biofilm biomass was quantified by measuring the optical density at 550 nm (OD_550_) using a SPECTROStar Nano microplate reader (BMG LabTech). Micrographs of stained biofilms were taken via microscopy as described above.

### RNA sequencing and analysis

Biofilms were cultured and dispersed as previously described with some adjustments ^19^. Briefly, tissue culture flasks (Nunc, ThermoFisher Scientific) were inoculated with 10^7^ CFU/ml cells in 50 ml of M9 medium from overnight cultures. Cultures were incubated at 37°C shaking at 70 rpm for 2, 4 or 8 h. At 2 h and 8 h, planktonic cells (10^8^) were collected from the liquid phase and immediately mixed with 2 volumes of RNA-protect solution (QIAGEN). Biofilm cells were gently rinsed with 10 ml of PBS and resuspended in 6 ml of RNAprotect (QIAGEN, Cat# 76506) mixed with 3ml of PBS using cell scrapers. Cells (10^8^) were subsequently pelleted at 5000 g (10 min, 25 °C). RNA was extracted with the RNeasy mini kit (QIAGEN, Cat# 74104) as per the manufacturer’s protocol for Gram negative bacteria grown in minimal media. Extracted RNA samples were treated with DNase I and rRNA was removed using streptavidin-coated magnetic beads (BGI Genomics, Shenzhen, China). Library was prepared by RNA fragmentation and cDNA synthesis followed by the ligation of adaptors at the 3’ end of cDNA fragments. These were further amplified by PCR and single-stranded fragments were circularised for downstream DNBSEQ PE100 sequencing (BGI Genomics, Shenzhen, China). Clean reads were mapped to the *P. aeruginosa* PAO1 reference genome (GenBank Accession number AE004091.2) with Bowtie2 v2.4.5 ^63^. Geneious Prime v2024.0.7 was used to generate the read count matrix and to subsequently run DESeq2 v1.50.2 to compare the expression levels between sample groups (attachment, biofilm maturation and dispersal).

The Principal Component Analysis (PCA) plot was conducted exclusively as an exploratory visualisation to assess overall transcriptomic similarity within biological replicates and to examine separation between treatment groups. It was generated by extracting the reads per kilobase of transcript per million mapped reads (RPKM) of every gene at each one of the stages (Table S1). Then, genes with <|1| log_2_ fold-change transcription or *P*-value >0.01 for all groups were discarded, and the remaining differentially regulated genes were plotted using GraphPad Prism 10.4.1 (Graph Pad, USA).

### Gene sequence conservation analysis

A local database was generated with 1301 complete *P. aeruginosa* genomes (taxid: 287) were downloaded from GenBank, excluding atypical, metagenome-assembled genomes and genomes from large multi-isolate projects. Gene sequences were extracted from the annotated *P. aeruginosa* PAO1 reference genome AE004091.2. BLAST analyses were performed locally using the NCBI BLAST plugin for Geneious Prime v2024.0.7 against the local *P. aeruginosa* database. In-built Megablast analyses were configured with minimum E value of 10^-8^ and query cover of 75%.

### RT-qPCR

To validate RNA-seq data, total RNA was extracted from cells harvested from different biofilm stages as described above. cDNA was transcribed using 1 µg of total RNA using the SuperScript III first-strand synthesis kit (Invitrogen) as per the manufacturer’s protocol. cDNA samples were then used as templates for RT-qPCR conducted with the Quantinova SYBR Green master mix (Qiagen) as per the manufacturer’s protocol. The housekeeping gene *recA* was used as a control. Each sample was independently assayed twice with the gene-specific primers listed in Table S2. Relative gene expression was calculated using the 2^-ΔΔCT^ method ^64^. Transcriptional fold changes were plotted using GraphPad Prism.

### Construction of *lacZ* fusion reporter plasmids and **β**-galactosidase assays

The cloning vector carrying a promoterless *lacZ* was constructed via Gibson Assembly with a 3.2 kb fragment containing *lacZ* from *E. coli* K-12 MG1655, a 150 bp fragment containing the MCS of pBBR1-MCS5 and a 3.8 kb fragment containing a gentamicin resistance cassette and the pBBR1 origin of replication from pBBR1-MCS5, resulting in the reporter construct pBBR1-lacZ. Gene promoter regions were predicted using Sapphire.cnn.pseudomonas ^65^. The 5’ untranslated region (∼500 bp) of each gene/operon containing putative promoters were cloned into the MCS by BamHI and HindIII upstream of *lacZ*. Reporters were introduced into *P. aeruginosa* PAO1 via electroporation.

β-galactosidase-based reporter assays were performed as previously described ^66^. Briefly, 10^8^ cells were collected by centrifugation from biofilm phases at early biofilm maturation or biofilm dispersal stages and OD_600_ was recorded. Cells were lysed with 1mg/ml lysozyme in TE buffer (10 mM Tris-HCl, 1 mM EDTA). Cell lysates were resuspended in 1ml of Z-buffer (0.06 M Na_2_HPO_4_, 0.04 M NaH_2_PO_4_, 0.01 M KCl, 0.001 M MgSO_4,_ 0.05 M β-mercaptoethanol) and 160 µl were transferred to a 96-well plate in triplicate (Nunc, ThermoFisher Scientific). Then, 40 µl of an ONPG solution (4mg/ml) in phosphate buffer (0.06M Na_2_HPO_4_, 0.04 M NaH_2_PO_4_; pH 7.0) were added to each well and reactions were incubated for 2 h at 28 °C in a plate reader. Absorbances were recorded at 420nm and 550nm every 10 min and Miller units (MU) were calculated using the following equation:

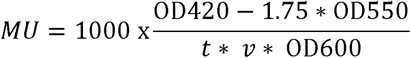

Where *t* is the reaction time, *v* is the volume of culture used and OD600 is the density of the culture prior to centrifuging.

Statistical differences in enzymatic activity between cells collected at early biofilm maturation or biofilm dispersal were calculated using Student t-tests (**, P-value <0.01; ***, P-value <0.001; ****, P-value <0.0001). One representative experiment is shown from two independently performed assays.

## Acknowledgements

This work was funded by an Australian Research Council project grant (DP210101317), the Max Planck Queensland Centre on the Materials Science of Extracellular Matrices, and a QUT Amplify Scholarship provided by the Queensland University of Technology (Australia) to XB. The Ian Potter Foundation sponsored the CLARIOStar high-performance microplate reader (BMG, Australia). The funders had no role in study design, data collection and analysis, decision to publish, or preparation of the manuscript. The authors would like to thank Professor Robert EW Hancock (University of Columbia) for providing the *P. aeruginosa* PAO1 strain used in this study.

## Author contributions

JQ and MT conceptualised the project. XB, JQ and MT contributed to experimental design. XB conducted all experiments, and contributed to data collection, analysis and visualisation. XB, JQ and MT contributed to data interpretation. JQ, MT and KFS supervised the project. KFS and MT obtained the funding. XB wrote the manuscript draft, JQ substantially revised the manuscript, all authors edited the manuscript.

## Data availability

The RNA-Seq data have been deposited in the NCBI Short Read Archive (SRA) database with accession code PRJNA1291829.

