## Supplementary material for "Transcriptional profiling of *Pseudomonas aeruginosa* biofilm life cycle stages reveals dispersal-specific biomarkers": Fig S1, Fig S2, Table S2

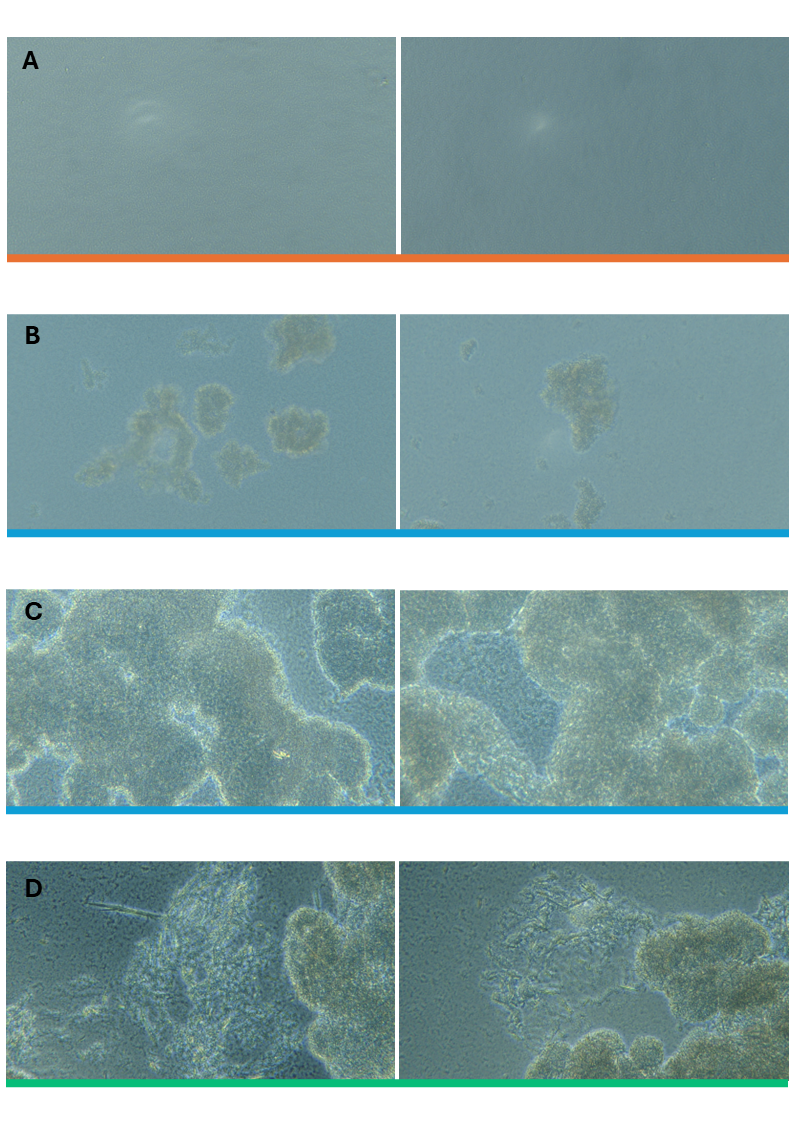

**Fig S1. Bright-field micrographs of *P. aeruginosa* PAO1 during each stage of the biofilm life cycle.** *P. aeruginosa* PAO1 cultures were seeded on 24-well plates and biofilm formation, and dispersal was recorded at different time points by bright-field microscopy at 400x. Images were captured from wells prior to crystal violet staining (Fig 1B). (A) Orange panels mark the early attachment phase (2 h); (B-C) blue panels mark the biofilm maturation stage (4 h and 8 h); (D) Green panels mark the dispersal stage (12 h).
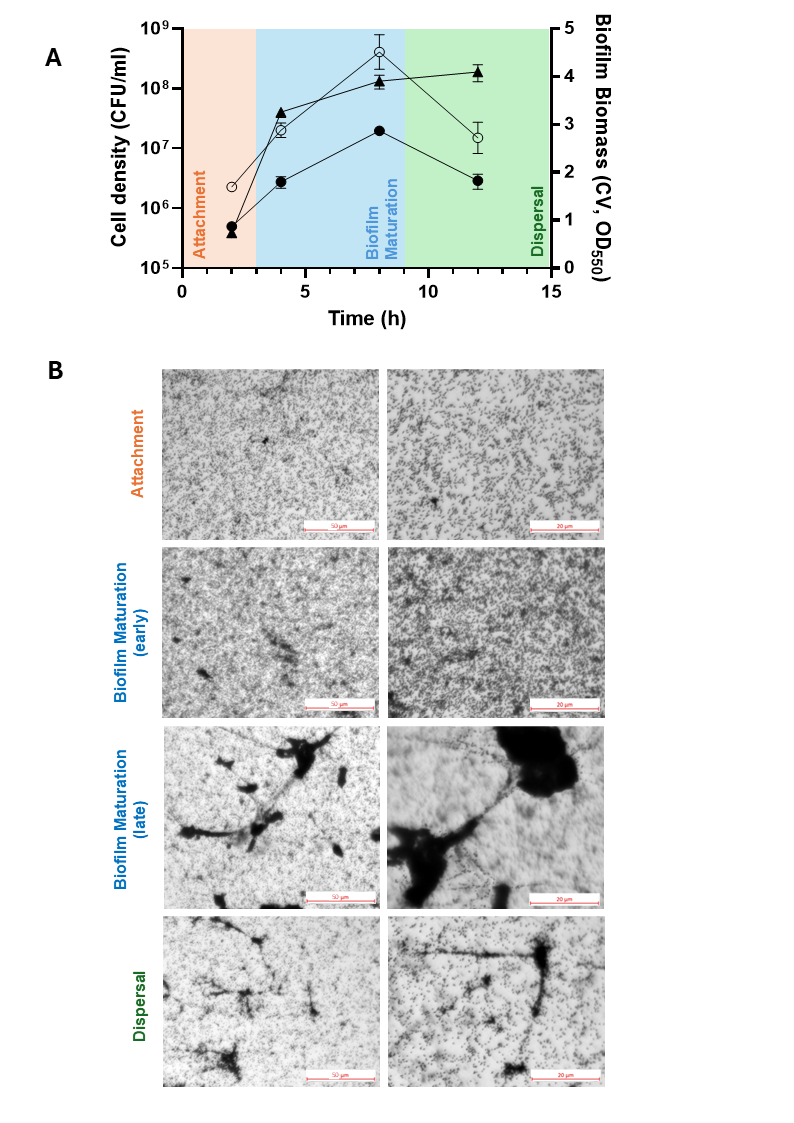

2 h

4 h

8 h

12 h

**Fig S2. Biofilm formation and dispersal kinetics of *P. aeruginosa* PAO1 in tissue culture flasks. (A)** *P. aeruginosa* PAO1 biofilm biomass and culture cell density over 12 h. PAO1 cultures were seeded on 24-well plates and biofilm biomass was quantified at different time points by crystal violet staining (CV, OD_550_; ○) and cell density (CFU/ml; ●). Culture density was simultaneously monitored (CFU/ml; ▲) with means ± SD shown. 3 biological replicates are represented. **(B)** Microscopic images of crystal violet-stained biofilms at each stage of the biofilm life cycle of *P. aeruginosa* biofilm cultures. Images are representative of at least 3 biological replicates. Orange: attachment stage. Blue: biofilm maturation stage. Green: dispersal stage. This colour coding scheme is used throughout the article to differentiate the three biofilm stages.

**Table S2: Primers used in this study**

| **Locus Tag** | **Direction** | **Sequence 5’ 🡪 3’** |
| --- | --- | --- |
| **RT-qPCR** | | |
| PA1985 F | Forward | GGT TTC GAA GTG ACC CTC TAC |
| PA1985 R | Reverse | CGA ACC GAG AAT ACG GAT GTG |
| PA4306 F | Forward | CCT GAC CCT GTT CGT GTA TT |
| PA4306 R | Reverse | TGG GAC TCA ATA CGG CAA TC |
| PA0743 F | Forward | AAC AGA TCG CCT TCA TCG G |
| PA0743 R | Reverse | CGA AGA CGT TCA GCA GGT AG |
| PA1353 F | Forward | CCT ACC TGA TCT TCA ACG GTA AT |
| PA1353 R | Reverse | GCG TGC ATG ATC TTG TCT TTC |
| PA4523 F | Forward | GAC GAC ACC CTC TTC GAA AT |
| PA4523 R | Reverse | TAG ACC CGC TCG ATA CTC TT |
| PA3385 F | Forward | ACA GGC AAC TCC TAC CTA CT |
| PA3385 R | Reverse | AGT TCA TGC TGC GGT GAT |
| PA0175 F | Forward | CAA CCA GGA GTT CCA CTA CAC |
| PA0175 R | Reverse | TAT AGA CCA GTT GCG CCT TG |
| PA4302 F | Forward | GAC CTT GCG CCT GAT GAT TT |
| PA4302 R | Reverse | GTC GAG ACT GAA CAG CGT ATT G |
| PA4304 F | Forward | CGG CGA CAT CAA GGT CAA T |
| PA4304 R | Reverse | CGC CGA AGA TCA GGT TGA AT |
| PA4648 F | Forward | CGG CAA TAT CCA GAT CCA GTG |
| PA4648 R | Reverse | GGC GTC CGA ATA GAT GTT GTA G |
| PA4305 F | Forward | ATG TCG ACG TGA TGC TCT TC |
| PA4305 R | Reverse | CTC TCG CCG TAG GTC AGT A |
| PA4649 F | Forward | GTA CGC CTC CCT GAA CAA C |
| PA4649 R | Reverse | GTC GAG CGT GAT CGC ATA G |
| PA2588 F | Forward | CGG GCA TAC GAG GAG TAT TT |
| PA2588 R | Reverse | GTT GGC AGT GGC GAT CT |
| PA0111 F | Forward | GCG ATT CCG ATG CCT GAA |
| PA0111 R | Reverse | CGG AAC CCA GAA ATT CCA TTT G |
| recA F | Forward | CAA CTG CCT GGT CAT CTT CAT C |
| recA R | Reverse | CGT AGA ACT TCA GTG CGT TAC C |
| rhlA F | Forward | CGAGACCGTCGGCAAATAC |
| rhlA R | Reverse | GCA CCT GGT CGA TGT GAA A |
| rhlB F | Forward | TGT CAC AAC CGC ACA GTA TC |
| rhlB R | Reverse | CTT CAG CCA TCG AGC ATC C |
| rhlC F | Forward | GTG CTG GTG GTA CTG TTC AA |
| rhlC R | Reverse | GTT GTC GAC GGC AAG GAA |
| dspS F | Forward | TGG TCA ACG AGC AAC AGG |
| dspS R | Reverse | CAA GCC ATA CTG GTA GAG TTC C |
| dspI F | Forward | CGC TGA TCA CCA TCA ACC A |
| dspI R | Reverse | GTC ACT ACC AGG GCG TAG ATA |
| eddA F | Forward | GCC GAC CAG TCG ATC TTC TA |
| eddA R | Reverse | GCT CCA GAC GAA ACG GAT ATT G |
| eddB F | Forward | CTA CCG CGT CGA GTT CTA TTT |
| eddB R | Reverse | GCG AGG ACG AAG GTC TTG |
| amrZ F | Forward | GAA ACA GGC AAC TCC TAC CTA C |
| amrZ R | Reverse | GAG CGA CTT CTG CGA TCT G |
| PA4781 F | Forward | TGA TCC TGC TCG ATG TGA AC |
| PA4781 R | Reverse | GGG CAG TGA GGA ACA TCA A |
| PA4108 F | Forward | GTA TGT TCG TCC AGA GCC TTT |
| PA4108 R | Reverse | ATC GAT CCA GAC CTC CTT CA |
| **Cloning primers** | | |
| PA0110 promoter F - BamHI | Forward | AAA AGG ATC CGC TCG ACC TTC TTC ATG C |
| PA0110 promoter R - HindIII | Reverse | AAT TAA GCT TGA GTC CCA CCA TGT TGA AAG |
| PA0179 promoter F - BamHI | Forward | AAA AGG ATC CGA TTC CAA GCA CAA GAT CC |
| PA0179 promoter R - HindIII | Reverse | AAA AAA GCT TAA TCG GTT TGC CCA TGG T |
| PA0743 promoter F - BamHI | Forward | AAT TGG ATC CCT TCA CAG CCT GCT TCG C |
| PA0743 promoter R - HindIII | Reverse | AAT TAA GCT TGG AGC TCC TCC TAT GGT G |
| PA1353 promoter F - BamHI | Forward | AAT TGG ATC CGC TAT CGT CTT GGA GCG G |
| PA1353 promoter R - HindIII | Reverse | AAT TAA GCT TCT GCG TTC TCC TTG TTG TTC |
| PA2588 promoter F - BamHI | Forward | AAT TGG ATC CAA GAT CGA GTT GAC CCT GC |
| PA2588 promoter R - HindIII | Reverse | AAT TAA GCT TCG GTT ACC TCT TGC ACG C |
| amrZ promoter F - BamHI | Forward | AAT TGG ATC CGA GCG TGG ATT TGC CGG |
| amrZ promoter R - HindIII | Reverse | AAT TAA GCT TCA TTG AAC CTG TAG AGT CAG G |
| flp promoter F - BamHI | Forward | AAT TGG ATC CGA GGT ATC CGA GCA ACA CC |
| flp promoter R - HindIII | Reverse | AAT TAA GCT TTC TTG TTT GCT CCT CGA ACG |
| PA4305 promoter F - BamHI | Forward | AAA AGG ATC CCC ACC GCG ATC AAA CCG |
| PA4305 promoter R - HindIII | Reverse | AAA AAA GCT TCA CCT TGC TAT TCA TGG GC |
| PA4523 promoter F - BamHI | Forward | AAT TGG ATC CCA GGT CAC CAG CAC TCG |
| PA4523 promoter R - HindIII | Reverse | AAT TAA GCT TCG GGA CTC TCC TGC CA |
| PA4648 promoter F - BamHI | Forward | AAA AGG ATC CGG TGA CCC TGG TGT TCG |
| PA4648 promoter R - HindIII | Reverse | AAA AAA GCT TGC GTT TAT TCA TCT CTA ACT TCC |
| PA1985 promoter F - BamHI | Forward | AAA AGG ATC CCT CAA GGG GCA CAA CGG |
| PA1985 promoter R - HindIII | Reverse | AAA AAA GCT TGA AGC TGG GCT TGG TCC |
| pBBR1-MCS5 (MCS) F | Forward | GTT CGT GCC TTC ATC CGC TCT AGA ACT AGT GGA TC |
| pBBR1-MCS5 (MCS) R | Reverse | CGT AAT CAT GGT CAT AAA GGG AAC AAA AGC TGG |
| E. coli K-12 MG1655 lacZ F | Forward | GCT TTT GTT CCC TTT ATG ACC ATG ATT ACG GAT TC |
| E. coli K-12 MG1655 lacZ R | Reverse | ACG GTC ACA CTG CTT TTA TTT TTG ACA CCA GAC C |
| pBBR1-MCS5 (backbone) F | Forward | TGG TGT CAA AAA TAA AAG CAG TGT GAC CGT GTG |
| pBBR1-MCS5 (backbone) R | Reverse | ACT AGT TCT AGA GCG GAT GAA GGC ACG AAC CCA G |
